# MultiCPA: Multimodal Compositional Perturbation Autoencoder

**DOI:** 10.1101/2022.07.08.499049

**Authors:** Kemal Inecik, Andreas Uhlmann, Mohammad Lotfollahi, Fabian Theis

## Abstract

Single-cell multimodal profiling provides a high-resolution view of cellular information. Recently, multimodal profiling approaches have been coupled with CRISPR technologies to perform pooled screens of single or combinatorial perturbations. This opens the possibility of exploring the massive space of combinatorial perturbations and their regulatory effects computationally from the extrapolation of a few experimentally feasible combinations. Here, we propose *MultiCPA*, an end-to-end generative architecture to predict multimodal perturbation response at single cell level. Two mixing strategies to integrate multiple modalities are introduced and compared with existing methods. MultiCPA was also shown to accurately predict unseen combinatorial perturbation responses for multiple modalities. The code to reproduce the results is available on *GitHub*, theislab/multicpa.

## 1. Introduction

Single-cell multiomics (Teichmann & Efremova, 2020) datasets are routinely generated to capture the cellular heterogeneity (Stephenson et al., 2021; Yao et al., 2021) with higher resolution through simultaneous quantification of transcriptome and surface proteins in the same cell (Stoeckius et al., 2017). Recently, multimodal technologies have been combined with CRISPR-compatible cellular indexing of transcriptomes and epitopes to profile cells under single (Mimitou et al., 2019; Frangieh et al., 2021) or combinatorial (Wessels et al., 2022) genetic perturbation. However, a comprehensive experimental investigation of combinatorial perturbations is challenging due to massive exploration space of possible combinations. Thus, computational methods are required to *in silico* predict cellular responses to a perturbation to navigate the large perturbation space and facilitate the experimental design.

Computational models based on representation learning (Lotfollahi et al., 2019; Ji et al., 2021) have been successfully applied to predict gene expression response to disease and chemical perturbation at the single-cell level. Recently, compositional perturbation autoencoder (CPA) (Lotfollahi et al., 2021) was proposed to predict gene expression response to combinatorial drug or genetic perturbations. Yet, CPA predictions are limited to a single modality, the gene expression. However, perturbation response prediction across multiple modalities helps to obtain a more holistic view of cellular behavior. Existing approaches such as Total Variational Inference (totalVI) (Gayoso et al., 2021) have been shown to efficiently model CITE-seq data by modeling biological and technical factors in the data. totalVI has been applied to perform counterfactual prediction to impute unmeasured surface proteins for single-cell RNA-seq data. However, the model is unable predict the combinatorial perturbation responses.

To address these challenges, we present multimodal compositional perturbation autoencoder, *MultiCPA*, an end-to-end generative model to exploit paired measurement of RNA and surface proteins to learn perturbation responses across both modalities at single-cell level. We demonstrate MultiCPA can efficiently model highly multiplexed multi-modal CRISPR screens to predict unseen single and combinatorial perturbations. Furthermore, MultiCPA learns a probabilistic representation of the data while accounting for biological and technical factors.

## 2. Methods

### 2.1. Integrating multimodal perturbation profiles

To evaluate the relative successes of two different mixture models, Product-of-Expert (PoE) (Lee & van der Schaar, 2021) and concatenation, two variational autoencoders (VAEs) (Kingma & Welling, 2013) were built as shown in Figure 1. In concatenation based model architecture, MultiCPA (concat), joint feature vectors are constructed by concatenating observed data from both modalities, proteins *(x*_*P*_) and genes (*x*_*G*_). Joint embedding, *Z*_*joint*_, is sampled via the reparameterization trick (Kingma et al., 2015) from joint posterior, *q(Z*_*joint*_ | *x*_*G*_,*x*_*P*_), which is estimated by a shared encoder from the concatenated feature vector. The PoE mixture model architecture, MultiCPA (PoE), estimates independent marginal latent distributions *q(Z*_*P*_ | *x*_*P*_) and *q(Z*_*G*_ | *x*_*G*_) by the respective encoder of each modality. Joint embedding, *Z*_*joint*_, is estimated similarly from joint posterior *q(Z*_*joint*_ | *x*_*G*_,*x*_*P*_), the product of the conditional marginal posteriors calculated using PoE framework. The loss function of the mixture module *ℒ*_M,1_ for MultiCPA (concat) is given in Equation 1, while arithmetic mean of all KL divergences is used for MultiCPA (PoE).

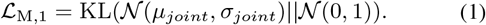

**Figure 1.**
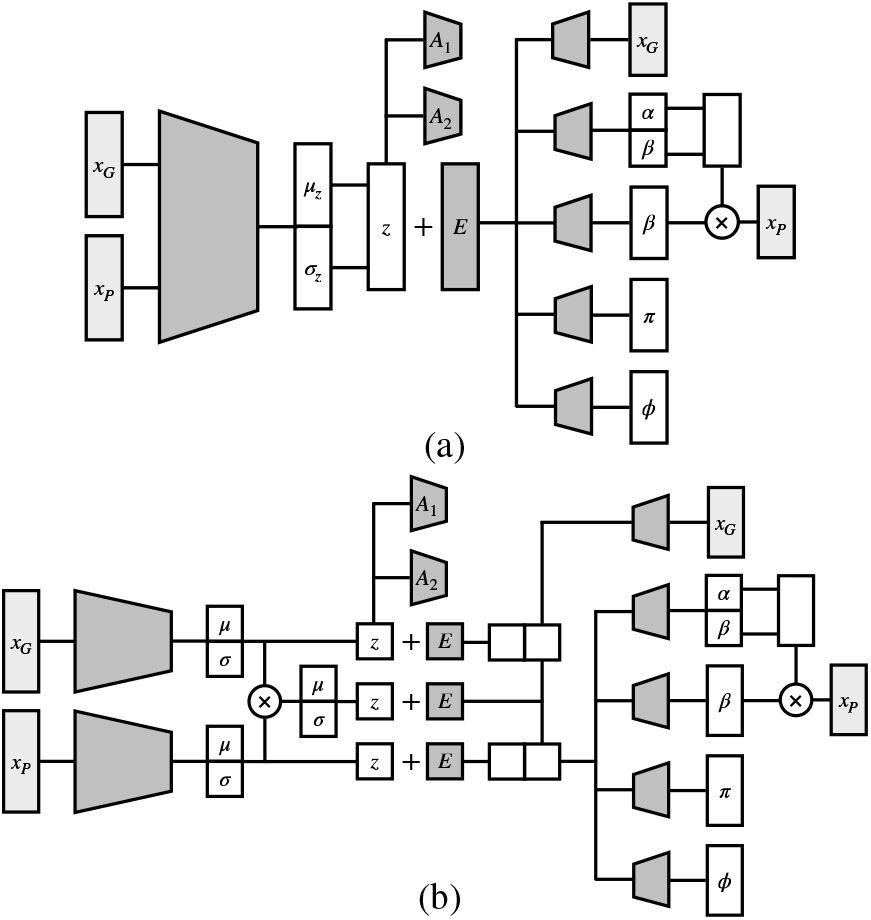
Overview of proposed MultiCPA architectures, where *A*_*i*_ denotes adversarial discriminator networks, *E* denotes separate perturbation and covariate embeddings. Multimodal integration by a) concatenation mixture module. b) PoE mixture module.

The information about perturbations and covariates in the joint embedding is disentangled by using an adversarial network as implemented in CPA framework (Lotfollahi et al., 2021). Auxiliary cross entropy losses implemented for two adversarial discriminator networks, which are trained to predict perturbation and cell covariates from *Z*_*joint*_, forces the encoders to produce the basal cellular state, *Z*_*basal*_. The adversarial loss for MultiCPA (concat) is given in Equation 2, while the adversarial loss for MultiCPA (PoE) is defined by the arithmetic mean of *ℒ*_*A*_*(Z*_*P*_), *ℒ*_*A*_*(Z*_*G*_) and *ℒ*_*A*_*(Z*_*joint*_).

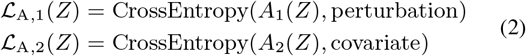

*Z*_*basal*_ is then composed with each of perturbation and covariate embeddings separately in the latent space and forwarded to the decoder networks. To reconstruct the observed data from the latent representation of the observations, the conditional latent space embeddings are entailed to learn perturbations and covariates. Extracting expressions (gene and protein), perturbations, and covariates as disentangled embeddings in the latent space allows to predict counterfactual scenarios such as unseen perturbation combinations.

### 2.2. Modality-specific data reconstruction

Joint latent space embedding feeds into the modality-specific decoders trying to reconstruct the corresponding input data. Inspired by totalVI concepts (Gayoso et al., 2021) integrating genes and protein modalities, the decoder network in MultiCPA architectures consists of five neural networks. All encoders, decoders, and adversarials of both MultiCPA models were built from fully-connected blocks. The gene data is decoded using a single decoder to reconstruct the observed data utilizing negative binomial loss function (Equation 3, indices are dropped for readability), where the distribution is specified by the mean *μ*_*G*_ and the inverse dispersion *θ*_*G*_.

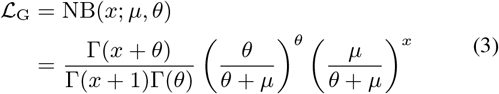

On the other hand, four separate decoders for background mean *μ*_*b*_, foreground mean *μ*_*f*_, protein mixing *π*_*P*_ of background and foreground, and protein dispersion *θ*_*P*_ are used to decode protein data. Negative binomial mixture loss comparing the decoded protein signal components with observed protein data was implemented to guide reconstruction procedure (Equation 4, indices are dropped for readability). Additionally, a KL divergence term, which utilizes a protein specific prior for the background mean learned during training, is computed.

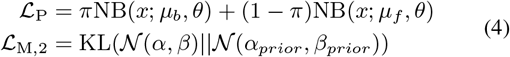

Decoding procedure during training is very similar for both models, except that MultiCPA (PoE) uses multiple respective latent space embedding instead. Optimization of the models during model training process repeats two successive steps. A batch from observed data passes through encoder networks to compute joint embedding *Z*_*joint*_, which feeds into perturbation and covariate discriminator adversarial networks. The stability of adversarial networks are improved by the implementation of gradient penalty to prevent gradients with large norm values. In the next iteration, joint embedding *Z*_*joint*_ is combined with perturbation and covariate embeddings in the latent space and then decoded through multiple decoders. Here, the total loss for back-propagation is given by Equation 5, where *w*_*i*_ are model hyperparameters.

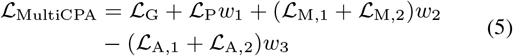

### 2.3. Hyperparameter tuning and datasets

Best hyperparameters of both models were extensively searched in a large hyperparameter space using the experiment management tool *sacred* (Greff et al., 2017) and a *MongoDB* (Gyorodi et al., 2015) experiment database on a large computer cluster with iterative sweeps. A certain set of hyperparameters were selected for each dataset and model combination, which optimize the dataset-specific counterfactual prediction accuracy in later analyses. Best models after hyperparameter tuning were manually examined in terms of reconstruction and adversarial losses in *train, test* and *OOD* (out-of-distribution) splits for both datasets to identify any possible overfitting issues. Two CITE-seq datasets of THP-1 human monocytic cells were used for model training and subsequent analyses. First dataset, named as *Wessels2022* (Wessels et al., 2022) has 30707 cells with 16920 genes and 24 proteins for 28 single gene knock-out perturbations. Second dataset, named as *Papalexi2021*, (Papalexi et al., 2021) has 20729 cells with 18649 genes and 4 proteins for 26 single gene knock-out perturbations, but no double perturbation. The datasets were quality checked, visualized and preprocessed using *scanpy* (Wolf et al., 2018). 5000 highly variable genes (HVGs) were selected for training the models. For each perturbation in each dataset, 20 differentially expressed (DE) genes were calculated to assess the model performances at test time.

## 3. Results

### 3.1. Choosing the best model architecture

Two VAE architectures with alternative mixture models were compared on the *Papalexi2021* dataset. Approximately 5000 models for each of the proposed architectures were trained on a high performance computing cluster to find best hyperparameter combination. The best models were chosen in terms of counterfactual perturbation prediction accuracy. Here, coefficient of determination, *R*^2^, is determined as the metric when comparing observed data with model predictions. The results in Figure 2a show that MultiCPA model with concatenation mixture module outperformed PoE based model in predicting protein data (0.98 vs. 0.57), although the reconstruction and adversarial losses were comparable.

**Figure 2.**
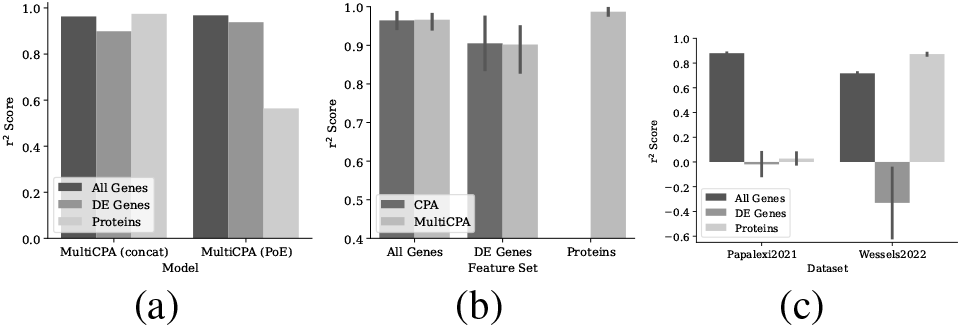
a) Prediction performances of the best models after hyperparameter tuning for the two MultiCPA models on *test* split. Comparison of MultiCPA and CPA in terms of *OOD* split prediction. c) Baseline expectation under random model behaviour.

MultiCPA (concat) was hence chosen as the best model in terms of total accuracy in counterfactual prediction, and MultiCPA (PoE) was omitted from subsequent analyses.

### 3.2. MultiCPA outperforms CPA leveraging additional modalities

MultiCPA was compared with CPA model in terms of the prediction accuracy on out-of-distribution (*OOD*) split. Both models were thus trained with *Wessels2022* dataset, where six perturbation combinations had been completely removed from the dataset and labelled as *OOD*. Figure 2b shows that MultiCPA model predicts counterfactual gene expression with slightly higher accuracy (0.96 ± 0.02 vs. 0.95 ± 0.02). Nevertheless, for all genes and differentially expressed (DE) genes in the dataset, both CPA and MultiCPA performed robustly (0.89 ± 0.07 vs. 0.88 ± 0.05). This suggests both MultiCPA and CPA not only extract and learn the individual effects of perturbations from *train* split, but also successfully combine these information in the latent space to predict the effect of unseen perturbation combinations. Additionally, MultiCPA predicts protein data with a very high accuracy for unseen perturbation combinations, leading to position itself as a multimodal extension of CPA.

Perturbation latent space in both datasets was inspected in order to assess whether perturbation embeddings could give clues about the similarities of perturbations’ mode of action. CPA has been previously shown to be competent in grouping perturbations together which are associated via cellular regulatory, metabolic or signaling pathways. It was hypothesized that additional protein information integrated with the MultiCPA model should result in a better resolution and biologically more meaningful groupings in perturbation latent space. Based on manual annotation of perturbations considering underlying biological mechanisms, MultiCPA and also CPA were not able to group the perturbations in both datasets in a consistent fashion, suggesting a dataset-specific variation or potential issues that still need to be accounted for (Figure 3). On the other hand, both models could successfully extract perturbation information from the input datasets although the datapoints of each perturbation do not cluster together on input feature set based UMAPs.

**Figure 3.**
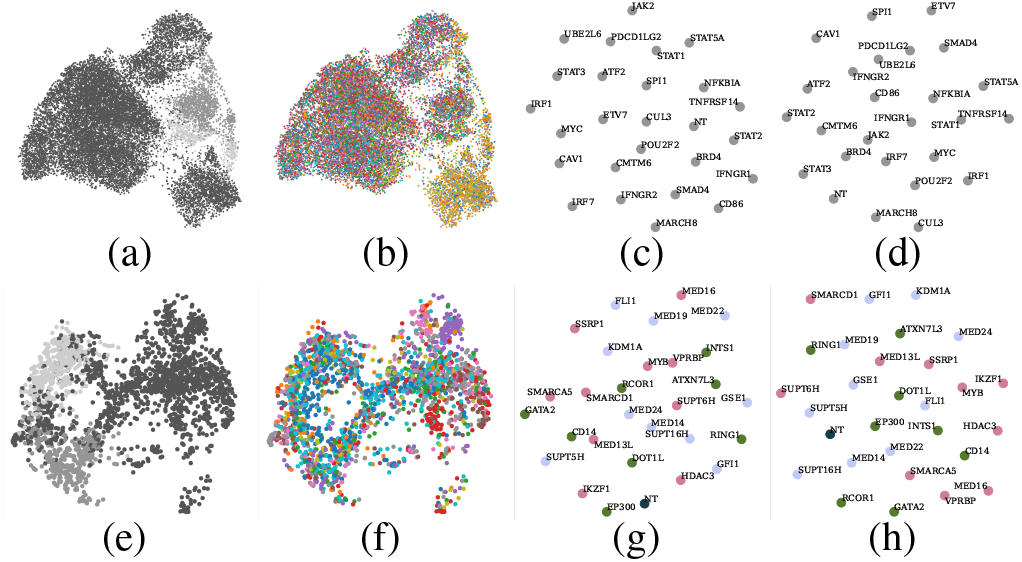
UMAP visualizations of input datasets and perturbation embeddings. First row is for *Papalexi2021*, second row is for *Wessels2022* for which only single perturbations are plotted for easier visual comparison. a, e) Input dataset UMAPs, where colors represent division phases G1, G2M, S, from darkest to lightest. b, f) Input dataset UMAPs, where colors represent single perturbations. c, g) Perturbation embeddings UMAPs for MultiCPA. d, h) Perturbation embeddings UMAPs for CPA.

### 3.3. Comparison with existing deep learning models

MultiCPA and totalVI are compared to assess their relative advantages in integration of a multimodal single-cell datasets to make counterfactual predictions. Learning the effect of individual perturbations from the input dataset and combining learned information to make a counterfactual combinational perturbation prediction is not possible using totalVI. The comparative analysis is thus conducted on *test* split only, but not *OOD* split. Considering each perturbation as a different batch, *batch transformation* method of totalVI was applied to unperturbed cells in the *test* set into the perturbation of interest to obtain model predictions of perturbation effects. Analogously, MultiCPA predictions were made only using the unperturbed cells. It was observed that MultiCPA compares slightly favorably to the existing totalVI method (Figure 4a and 4b).

**Figure 4.**
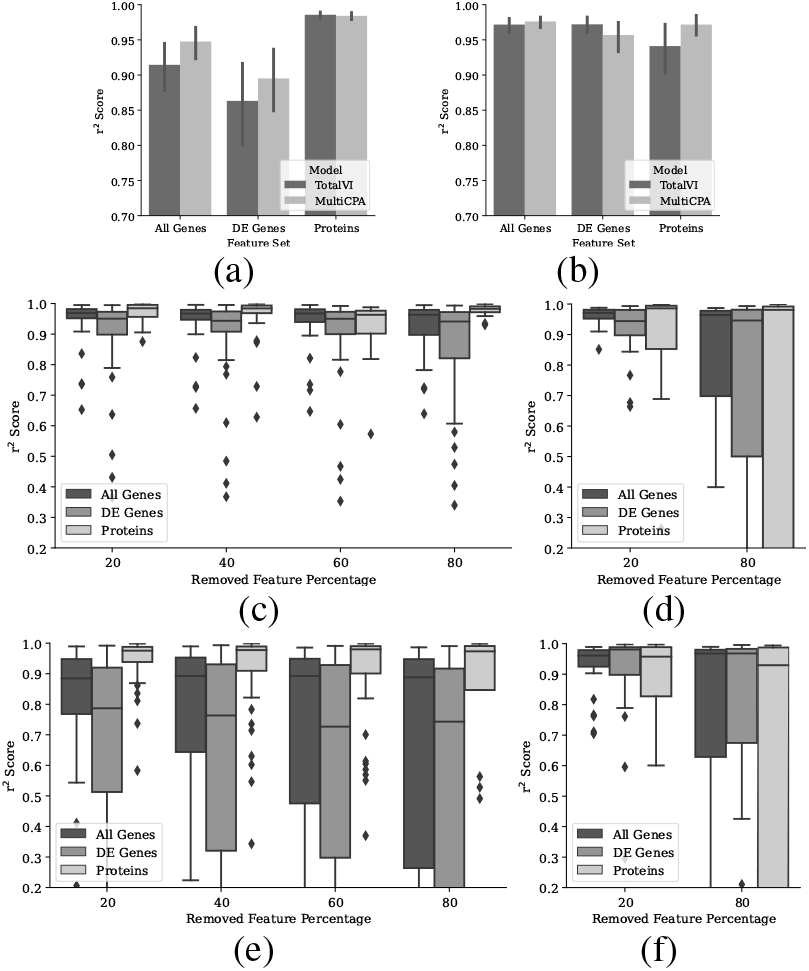
Counterfactual perturbation prediction performances on *test* split, respectively for *Wessels2022* and *Papalexi2021* datasets at each row. a, b) Prediction performances of totalVI and MultiCPA models. c, d) Prediction performances of MultiCPA model with noise added datasets. e, f) Prediction performances of totalVI model with noise added datasets.

With the idea of testing the robustness of the model to the noise found in the data, certain percentages of the input features were randomly selected from a random quarter of the perturbations in both datasets, and were replaced with zero values. Both totalVI and MultiCPA models were then trained with modified datasets using the same hyper-parameters tuned for the complete datasets, and counterfactual prediction accuracies were calculated as usual. totalVI model was considerably affected by such an intervention while MultiCPA still retains high prediction accuracies even with the most severe scenario for *Wessels2022* dataset (Figure 4). The responds to the intervention were comparable for *Papalexi2021* dataset. These results suggest that the information of each perturbation effect could be learned via untouched perturbations for MultiCPA but not totalVI, which is then used in test time to predict intervened perturbations in the dataset, as *Wessels2022* dataset contains many combined perturbations. However, *Papalexi2021* dataset contains only single perturbations, making it unlikely to learn intervened perturbations for both models.

## 4. Discussion

Harnessing the strength of multiple modalities in extracting the information regarding cellular effect of perturbations helps to acquire a broader insight into perturbation effects. Here, we showed that MultiCPA can provide a more comprehensive characterization of cellular phenotypes and perturbations in an unbiased manner through a joint multimodal data representation and improves the prediction of unseen perturbation combinations with higher accuracy. Moreover, MultiCPA is the first method that learns the effect of individual perturbations on the surface protein data and uses the information to predict unseen perturbation combinations.

Furthermore, the proposed generative deep learning model outperformes the existing totalVI model for multimodal single-cell data integration with regards to overall prediction accuracy in the *test* split. MultiCPA learns individual perturbation and covariate information from combinations and performs more robustly than totalVI in response to the noise in high-dimensional input data. While totalVI is not devised to learn and combine individual perturbations, MultiCPA exploits learned latent space embeddings to predict unseen combinations in the training data for both modalities. Additionally, relative advantages of two mixture models has been tested in this context, where MultiCPA (concat) was shown to outperform MultiCPA (PoE) in terms of counterfactual prediction accuracy of surface protein data.

We anticipate MultiCPA to guide exploration of the perturbation space, leading to the development of novel therapeutics (Brochado et al., 2018), and facilitating the discovery of the general principles of a cellular machinery (Muscato et al., 2022) in biological research. As future directions, we aim to extend our model with other cellular modalities, such as chromatin accessibility data by scATAC-seq (Lareau et al., 2019; Lotfollahi et al., 2022) technology.

